# Production, Purification, and Crystallization of Recombinant HER2 Tyrosine Kinase Domain (HER2-TKD) as a Platform for Structure-Based Drug Screening

**DOI:** 10.1101/2025.06.02.657378

**Authors:** Edanur Topalan, Halilibrahim Ciftci, Hasan DeMirci

## Abstract

The human epidermal growth factor receptor 2 tyrosine kinase domain (HER2-TKD) plays a central role in signal transduction and is a significant therapeutic target in cancer. This study aimed to produce soluble recombinant HER2-TKD in Escherichia coli to enable structural studies for drug screening applications. The HER2-TKD gene was cloned into the pET28a(+) expression vector and expressed in E. coli. Initial expression led to the formation of inclusion bodies; thus, sarcosyl was used to solubilize the aggregated protein. Several induction durations were tested to optimize soluble expression. SDS-PAGE analysis was used to monitor expression and solubilization efficiency. The recombinant protein was purified using size-exclusion chromatography and reverse affinity chromatography to remove the SUMO tag. Crystallization trials were initiated using commercial screens to obtain diffraction-quality crystals. Soluble HER2-TKD was successfully obtained after optimization of induction and solubilization conditions. Crystallization efforts are ongoing to improve crystal quality for future structural analysis. These results provide a foundation for structure-based drug discovery studies targeting HER2.

## 1. Introduction

The epidermal growth factor receptor 2 (HER2), also known as ErbB2, is a member of the ErbB family of receptor tyrosine kinases (RTKs), which are key regulators of cellular proliferation, differentiation, and survival. HER2 plays a critical role in cancer biology, particularly in breast cancer and non-small cell lung cancer (NSCLC) [1]. In many cancers, mutations in the tyrosine kinase domain (TKD) of HER2 are frequently detected. These mutations often induce conformational alterations in the ATP-binding pocket (ATP-BP), resulting in increased kinase activity and activation of oncogenic signaling pathways [2,15,21,23,24].

Unlike other ErbB family members that require ligand binding for dimerization and activation, HER2 is constitutively active in a ligand-independent manner [3]. Its potent ability to form heterodimers with other ErbB receptors enhances its signaling capacity, making HER2 deregulation a significant driver of tumorigenesis [1]. In breast cancer, HER2 amplification is often the primary oncogenic event. In contrast, HER2-driven NSCLC involves more diverse mechanisms [22,29–31], including gene amplification, activating mutations [20,26,27], and overexpression [5,16–19].

Under physiological conditions, HER2 activity is tightly regulated by a balance between its phosphorylated (active) and dephosphorylated (inactive) forms, which is essential for normal cellular growth and breast tissue development [5]. However, when this balance is disrupted by overactivation or mutation, it leads to uncontrolled cell proliferation and survival [6–10]. Although HER2-targeted monoclonal antibodies (mAbs) and earlier-generation pan-HER tyrosine kinase inhibitors (TKIs) have shown limited efficacy in treating HER2-altered NSCLC, newer HER2-specific TKIs have demonstrated more encouraging results [3]. Nonetheless, resistance mechanisms—such as the activation of alternative RTK pathways and upregulation of downstream survival signals—remain a challenge for both TKIs and mAbs [4].

In this paper, we describe the successful expression of HER2-TKD in *Escherichia coli (E. coli)*, where the protein first forms inclusion bodies. The recombinant HER2-TKD was solubilized in %1.5 sarcosyl, purified using affinity and size-exclusion chromatography, and verified by SDS-PAGE. Crystallization investigations commenced utilizing commercially available sparse matrix screens to acquire diffraction-quality crystals. This paper outlines a reproducible and effective technique for the creation and purification of soluble HER2-TKD, making it appropriate for structural investigations. Creating this platform is crucial for enabling structure-based drug discovery targeting HER2, especially in cancers with abnormal HER2 activity.

## 2. Materials and Method

### 2.1 Preparation for Protein Expression

#### 2.1.1 Gene Construct Design

HER2-TKD: **MSDSEVNQEAKPEVKPEVKPETHINLKVSDGSSEIFFKIKKTTPLRRLMEA FAKRQGKEMDSLRFLYDGIRIQADQTPEDLDMEDNDIIEAHREQIGG**MSGAAPNQ ALLRILKETELRKVKVLGSGAFGTVYKGIWIPDGENVKIPVAIKVLRENTSPKANKEILDEAYVMAGVGSPYVSRLLGICLTSTVQLVTQLMPYGCLLDHVRENRGRLGSQDLLNWCMQIAKGMSYLEDVRLVHRDLAARNVLVKSPNHVKITDFGLARLLDIDETEYHADGGKVPIKWMALESILRRRFTHQSDVWSYGVTVWELMTFGAKPYDGIPAREIPDLLEKGERLPQPPICTIDVYMIMVKCWMIDSECRPRFRELVSEFSRMARDPQRFVVIQNEDLGPASPLDSTFYRSLLEDDDMGDLVDAEEYLVPQQGAAAS*

**Bold:** Sumo tag

We designed a construct encoding the tyrosine kinase domain of human HER2 for recombinant protein production in *E. coli*. The gene sequence was cloned into the *pET28a(+)* vector, which has a kanamycin resistance gene. To facilitate purification, we added an N-terminal 6xHis tag. Additionally, we fused a SUMO tag to enhance the solubility of the recombinant protein. After completing the vector construction, we proceeded with the transformation experiments.

#### 2.1.2 Transformation Procedure

We dissolved 2 μg of the HER2 plasmid in 25 μL of distilled water and vortexed the solution for 30 seconds. After centrifuging at 9,000 RPM for 4 minutes, we repeated the vortexing and centrifugation steps to ensure homogeneity. We stored the prepared plasmids at -20°C until use.

For transformation, we mixed 2 μL of the plasmid DNA with 50 μL of *E. coli* Rosetta-II competent cells that had been thawed on ice. We incubated the mixture on ice for 20 minutes, followed by a heat shock at 42°C for 45 seconds. Immediately after, we put the cells on ice for 2 minutes. We then added 500 μL of sterile LB medium and incubated the cells at 37°C for 90 minutes, gently inverting the tubes every 15 minutes. Following incubation, we spread 50 μL of the culture onto LB agar plates containing kanamycin (50 μg/mL) and chloramphenicol (35 μg/mL). The plates were incubated overnight at 37°C.

The next day, we selected individual colonies and inoculated them into 10 mL of LB broth, cultivating them overnight at 37°C. We confirmed successful growth by observing turbidity in the cultures. To prepare glycerol stocks for long-term storage, we mixed 750 μL of the overnight culture with 750 μL of sterile 80% glycerol in cryotubes. After vortexing -to ensure homogeneity-, we stored the glycerol stocks at -80°C.

### 2.2 Mini-Checks

#### 2.2.3 Mini-Scale Production of HER2 Protein and Incubation Time Optimization

Following the successful transformation, we proceeded to evaluate protein expression. A 10 mL volume of LB medium was sterilized by autoclaving, and once cooled to a temperature safe to touch, antibiotics were added at a 1:1000 ratio (10 μL kanamycin and 10 μL chloramphenicol).

Chloramphenicol was used due to the resistance conferred by the Rosetta-II strain, while kanamycin was added for the selection of the pET28a(+) plasmid. After the addition of antibiotics, a portion of the HER2 glycerol stock, stored at -80°C, was inoculated into the medium and incubated overnight at 37°C.

The next day, protein expression was induced by adding 0.4 M IPTG at a 1:1000 dilution (Final concentration of IPTG is 0.4 mM). During incubation at 37°C, 1 mL samples were collected at 1, 2, 3, 4, 5, 6, 7, 8, and 16 hours post-induction. The samples were centrifuged at 6000 RPM for 5 minutes to pellet the cells and separate them from the LB medium. Each cell pellet was resuspended in 750 μL of lysis buffer and subjected to sonication. Finally, 40 μL of each lysate was mixed with 10 μL of loading dye, prepared for gel analysis, and analyzed by 12% SDS-PAGE.

#### 2.2.3 Solubility Analysis of HER2 Protein in Mini-Scale Production

After obtaining the desired expression results, we proceeded to assess the solubility of the HER2 protein. The culture was grown in 10 mL LB medium at 37°C until an OD₆₀₀ of 0.6 was reached, at which point expression was induced with 0.4 mM IPTG and incubation continued for 7 hours. The resulting 10 mL culture was equally divided into two 5 mL aliquots and harvested by centrifugation.

To evaluate the solubility of HER2 and to assess the solubilizing effect of sodium lauroyl sarcosinate (sarcosyl) in case of formation of inclusion bodies, we designed an experiment comparing two distinct lysis buffers: one with sarcosyl and one without. For the sample labeled “C,” lysis was performed using a buffer containing 200 mM NaCl, 20 mM imidazole, 50 mM Tris-HCl, 5% glycerol, and 0.1% Triton X-100 (pH 8.5). For the sample labeled “S,” the lysis buffer was composed of 10 mM Tris, 150 mM NaCl, 1 mM PMSF, 5 mM BME, and 1.5% sarcosyl (pH 8.5).

After the addition of a lysis buffer, the samples were sonicated at 41% amplitude in three cycles of 15 seconds each. Following sonication, whole cell lysates (WCL) were collected for gel analysis. The samples were then centrifuged at 6000 RPM for 5 minutes, and both pellet and supernatant fractions were prepared for SDS-PAGE. All samples were analyzed on 12% SDS-PAGE gels.

### 2.3 Large Scale Protein Production

Following the optimization of incubation time and solubility conditions for HER2 protein, we proceeded to large-scale production. For this purpose, four 1 L LB cultures were grown at 37°C until an OD₆₀₀ of 0.6 was reached. Protein expression was induced with 0.4 mM IPTG, and incubation continued for 7 hours. At the end of the incubation period, the cultures were harvested by centrifugation at 3500 RPM for 45 minutes.

The resulting cell pellets were resuspended in lysis buffer (10 mM Tris, 150 mM NaCl, 1 mM PMSF, 5 mM BME, 1.5% sarcosyl) and subjected to sonication at 41% amplitude, using 11 cycles of 20 seconds each, until the viscosity was visibly reduced.

Following sonication, the lysate was clarified by ultracentrifugation at 35,000 RPM for 1 hour to separate the pellet and supernatant fractions. To remove any residual effects of the sarcosyl detergent, the supernatant was dialyzed against 2 × 1 L of dialysis buffer at 4°C overnight using a dialysis membrane. After dialysis, the sample was loaded onto a nickel-nitrilotriacetic acid (Ni-NTA) affinity column to initiate the purification process. Samples were collected and analyzed at each purification step.

The buffers used during purification were as follows:

**Histidine A Buffer:** 150 mM NaCl, 20 mM Tris-HCl, 5% glycerol, pH 8.0

**Histidine B Buffer:** 150 mM NaCl, 20 mM Tris-HCl, 250 mM imidazole, 5% glycerol, pH 8.0

#### 2.3.1 Purification and Tag Removal

To further purify the HER2 protein, size exclusion chromatography (SEC) was performed using a Superdex-200 column equilibrated with a buffer containing 150 mM NaCl and 20 mM Tris-HCl at pH 8.0. During elution, fractions of 5 mL each were collected based on increases in UV absorbance. Aliquots from each fraction were analyzed by SDS-PAGE.

The SEC chromatogram displayed two prominent peaks. SDS-PAGE analysis of the corresponding fractions revealed that the first peak contained a higher amount of HER2 protein. Fractions 9, 10, 11, 12, and 13, which showed intense bands corresponding to HER2, were pooled (**Figure 3**).

The pooled fractions were concentrated using 30 kDa Amicon centrifugal filters by centrifugation at 2500 RPM for 15 minutes, repeated three times. The concentrated protein sample was then prepared for tag removal.

For cleavage of the His₆-SUMO tag, the concentrated HER2 protein was incubated with ULP protease at a ratio of 5:1000 (w/w) in a Falcon tube at 4°C for 40 minutes. Following incubation, the sample was passed through a Ni-NTA column to separate the cleaved HER2 protein from the His₆-SUMO tag and uncleaved species.

The buffers used during this step were as follows:

**Histidine A Buffer:** 150 mM NaCl, 20 mM Tris-HCl, 5% glycerol, pH 8.0

**Histidine B Buffer:** 150 mM NaCl, 20 mM Tris-HCl, 250 mM imidazole, 5% glycerol, pH 8.0

At each step, flow-through and elution fractions were collected for analysis. Samples from each stage, along with pre-cleavage controls, were analyzed by SDS-PAGE to confirm the efficiency of tag removal and purification.

### 2.4 Crystallization

HER2 protein samples were prepared in various forms, including:

● SUMO tag-cleaved protein
● SUMO tag-uncleaved protein

Crystallization trials were set up using the microbatch crystallization method in Terasaki plates. Over 3000 crystallization conditions **(Table 1.)** were screened per trial, covering multiple screening sets. Crystallization experiments were conducted at both +4°C and room temperature. In total, six rounds of crystallization screening were carried out, including both apo HER2 and HER2-inhibitor complexes. Plates were monitored daily under a microscope to detect any crystal formation.

**Table 1.** Condition List. A list of the crystallization screening conditions used in this study.

|  |  |
| --- | --- |
| Natrix I | Wiz Synergy IV |
| Natrix II | Pact Premier I |
| Wizard I | Pact Premier II |
| Wizard II | PEG Ion I |
| Wizard II | PEG Ion II |
| Wizard IV | Salt RX I |
| Crystal Cryo I | Salt RX II |
| Crystal Cryo II | Proplex I |
| Crystal Screen I | Proplex II |
| Crystal Screen II | MembFAC |
| Crystal Screen II | Crystal Lite |
| Wizard Cryo I | PEG RX I |
| Wizard Cryo II | PEG RX II |
| Helix I | Multi XTAC |
| Helix II | Macro SOL |
| Midas I | 3D Structure Screen |
| Midas II | Stura Footprint |
| NR-LBD I | Clear Strategy Screen I & II (pH 4.5) |
| NR-LBD II | Clear Strategy Screen I & II (pH 5.5) |
| PGA-LM I | Clear Strategy Screen I & II (pH 6.5) |
| PGA-LM II | Clear Strategy Screen I & II (pH 7.5) |
| Morpheus I | Clear Strategy Screen I & II (pH 8.5) |
| Morpheus II | Quick Screen / Ionic Liquid Screen |
| Structure I | GRID/NaCl – Na Malonate (*) |
| Structure II | GRID/PEG 6000–Ammonium Sulfate(*) |
| Index I | GRID/MPD – PEG LiCl (*) |
| Index II | JENA BC Nuc Pro 1/2 |
| Wiz Synergy I | JENA BC Nuc Pro 3/4 |
| Wiz Synergy II | JCSG-I |
| Wiz Synergy III | JCSG-II |

## 3. Results

### 3.1 Overexpression and Purification of Soluble HER2-TKD

Soluble HER-TKD was successfully overexpressed, and the expression parameters were optimized for maximum yield. The best expression was obtained by inducing with 0.4 mM IPTG at 37°C and incubating for 7 hours **(Figure 1)**. Sarcosyl was used as a solubilizing agent to reduce inclusion body formation, and the concentration of 1.5% [14] **(Figure 2)**. Despite the presence of a 6xHis tag in the construct, purification using Ni-NTA affinity chromatography was inefficient, most likely due to poor binding or detergent interference. As a result, size exclusion chromatography (SEC) was chosen as an alternative purification strategy. Two peaks were observed in the SEC chromatogram, and SDS-PAGE analysis confirmed that HER2-TKD was present in the first peak **(Figure 3)**. Fractions 9 to 13 from SEC were pooled and concentrated using 30 kDa Amicon.

**Figure 1.**
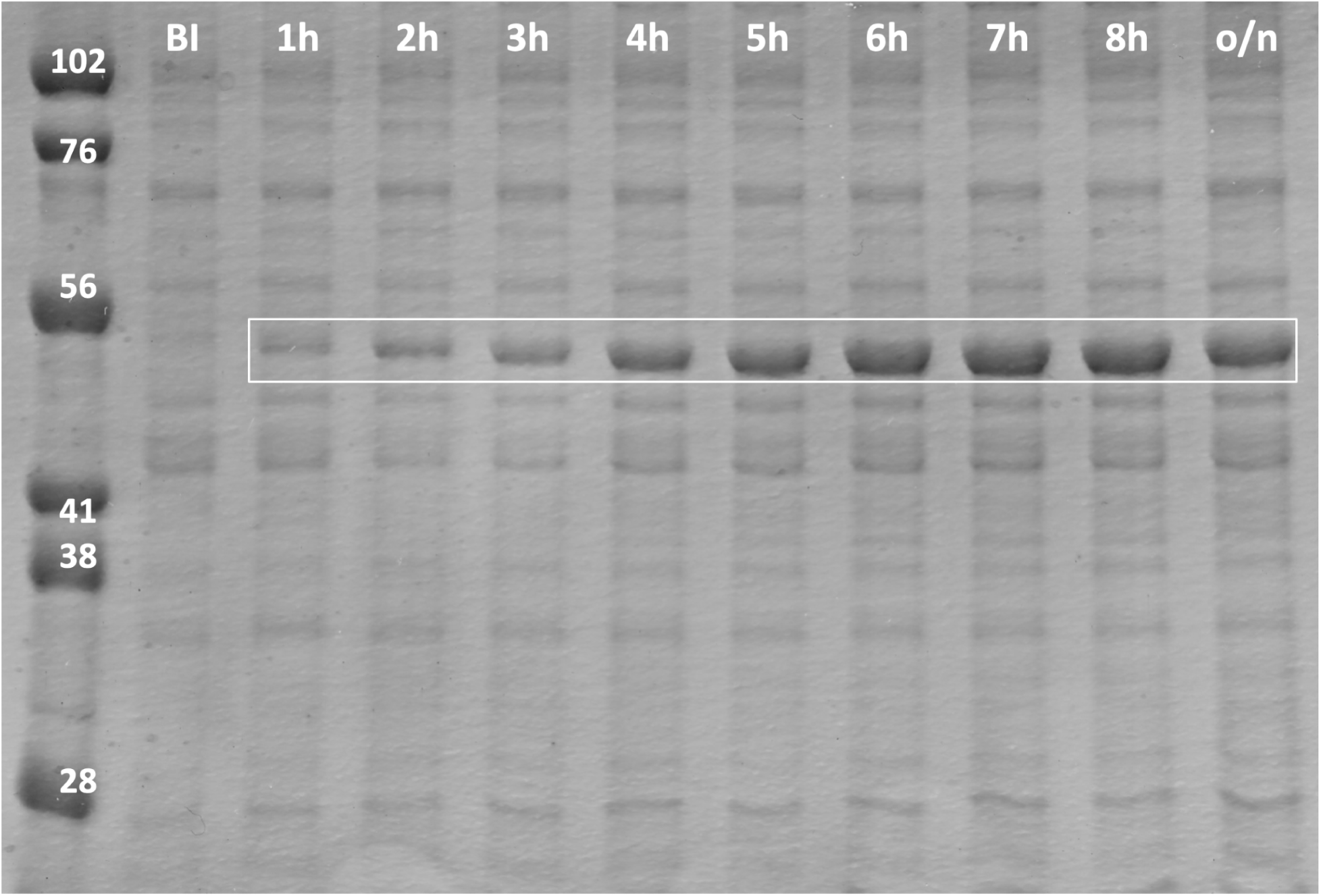
SDS-PAGE analysis of HER2 protein expression at different induction times. HER2 protein expression was evaluated after IPTG induction at various time points (1, 2, 3, 4, 5, 6, 7, 8, and overnight (o/n : 16 hours). Whole-cell lysates (WCL) corresponding to each induction time were loaded onto a 12% SDS-PAGE gel. Lane 1: protein marker; lane 2: cell lysate before induction (BI). Lanes 3 to 11 show samples collected at 1 h (Lane 3), 2 h (Lane 4), 3 h (Lane 5), 4 h (Lane 6), 5 h (Lane 7), 6 h (Lane 8), 7 h (Lane 9), 8 h (Lane 10), and 16 h (Lane 11) post-induction. Lane 11 represents the overnight induction sample. The prominent band corresponding to HER2-TKD (approximately 48 kDa) became more intense up to 7 hours of induction, after which no significant increase was observed. Based on these results, 7 hours of IPTG induction was determined to be optimal for efficient HER2-TKD production. A volume of 3 µL was loaded for all samples.

**Figure 2.**
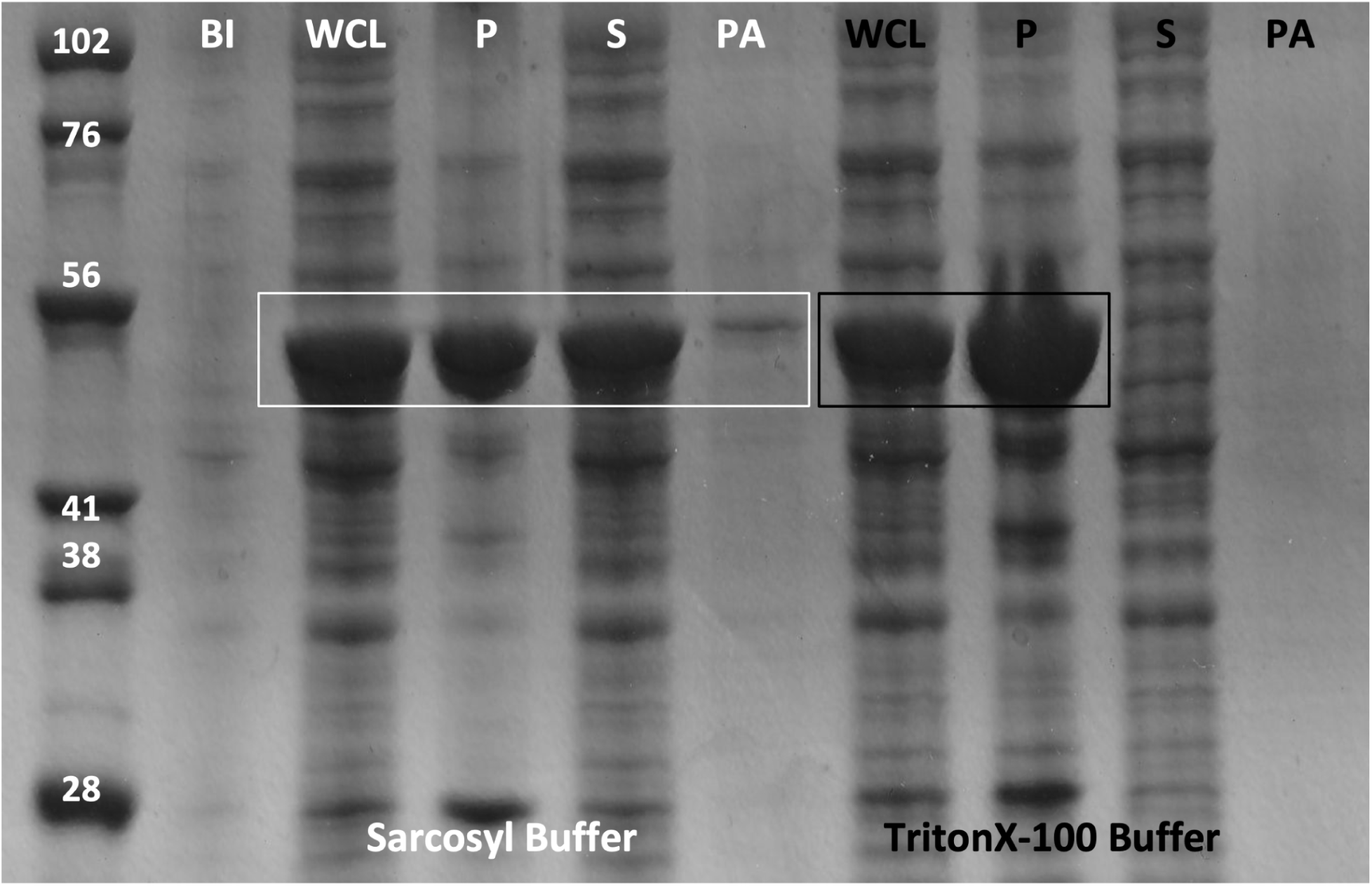
Solubility analysis of HER2 protein with and without Sarkosyl detergent by 12% SDS-PAGE. HER2 protein was expressed in *E. coli* and induced with 0.4 mM IPTG at 37°C for 7 hours. After harvesting, cells were lysed using buffer systems containing either 1.5% Sarkosyl (pH 8.5) or TrionX-100 (no Sarkosyl), to evaluate the detergent’s effect on protein solubility. Whole-cell lysates (WCL), soluble fractions (supernatants:S), and insoluble fractions (pellets:P) were separated by centrifugation and analyzed by 12% SDS-PAGE. Lane 1 contains the protein marker. Lanes 3–7 correspond to samples prepared in buffer with Sarkosyl: before induction (Lane 2), WCL (Lane 3), pellet (Lane 4), supernatant (Lane 5), and pull-down (PA) fraction (Lane 6). Lanes 8–11 show samples prepared with TritonX-100: WCL (Lane 7), pellet (Lane 8), supernatant (Lane 9), and pull-down fraction (Lane 10). The HER2-TKD band (∼48 kDa) was predominantly observed in the pellet fraction with TrionX-100, indicating poor solubility. In contrast, a significant portion of HER2-TKD shifted to the soluble fraction when Sarkosyl was included, demonstrating the detergent’s positive effect on solubilization. Based on these results, further HER2 protein purification was continued using Sarkosyl-containing buffers. A volume of 3 µL was loaded for all samples.

### 3.2 Tag Cleavage, Affinity Re-Purification, and Concentration of HER2-TKD

We performed HER2-TKD cleavage with ULP1 protease at an enzyme-to-substrate ratio of 5:1000 and incubated at 4°C for 30 minutes. This condition was chosen to maximize cleavage efficiency while maintaining protein stability and reducing degradation or aggregation, as demonstrated in our previous EGFR-TKD study. Following the cleavage, Ni-NTA affinity chromatography was used to separate the tags and the ULP1 protease, from the untagged HER2-TKD protein **(Figure 4).** The HER2-TKD flow-through concentrated using Amicon filters with a molecular weight cut-off of 30 kDa. However, protein solubility decreased after tag removal, and the HER2-TKD protein could not be concentrated above 2.5 mg/mL, posing a limitation for subsequent crystallization trials.

**Figure 3.**
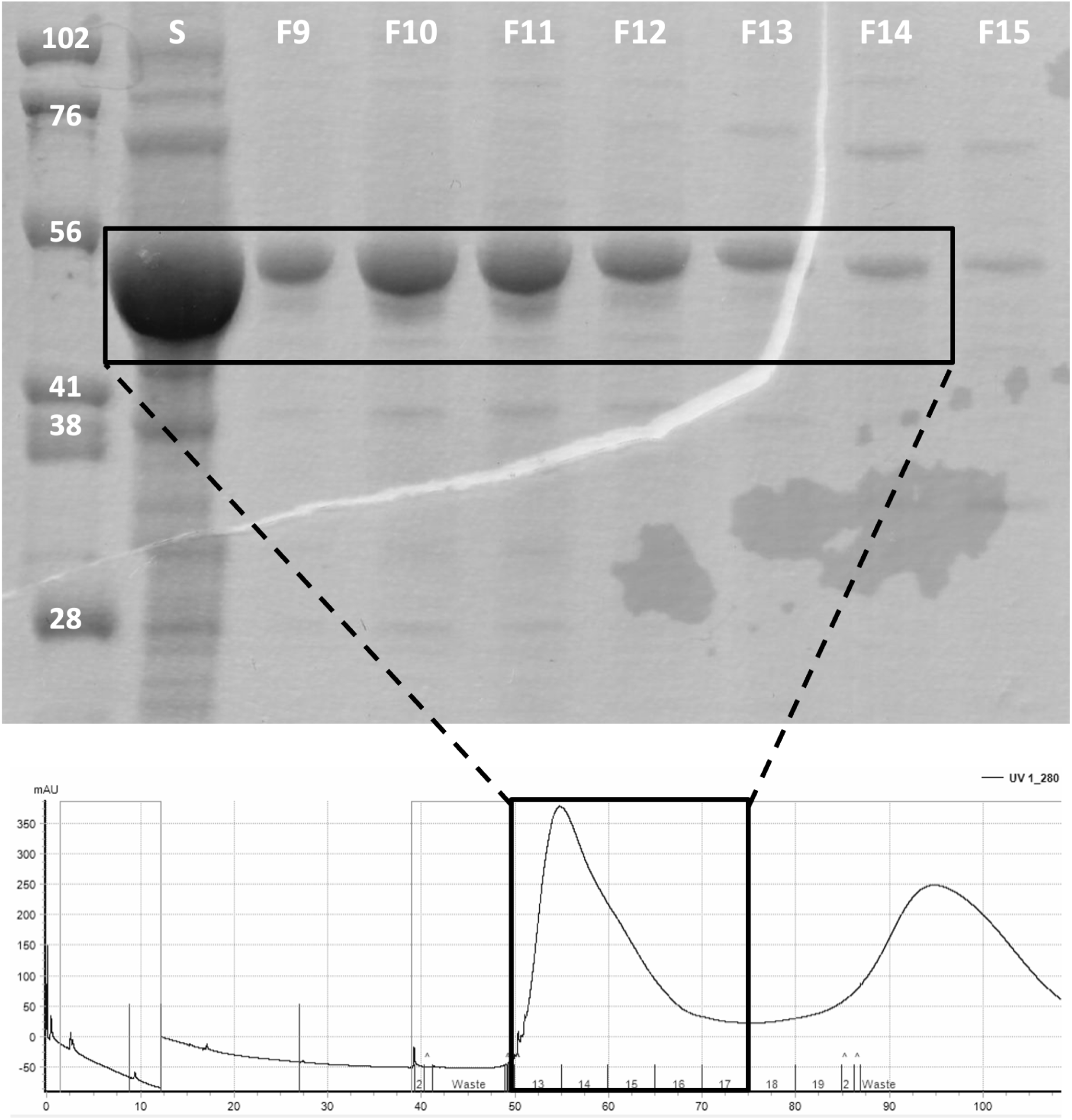
Size exclusion chromatography (SEC) purification of HER2-TKD and analysis of collected fractions by 12% SDS-PAGE. HER2-TKD protein was purified using SEC. In the chromatogram, fractions (F) 9–13 and 14–21 were collected as indicated by box. These fractions were analyzed by 12% SDS-PAGE. A clear band corresponding to HER2-TKD (∼48 kDa) was predominantly observed in fractions 9–13, suggesting that the target protein was enriched in this range. The lane assignments are as follows: Lane 1 contains the protein marker. Lanes 2 to 6 correspond to fractions 9 to 13, respectively. The results confirmed that fractions 9–13 contained the highest purity HER2-TKD protein and were selected for further experiments. A volume of 3 µL was loaded for all samples.

**Figure 4.**
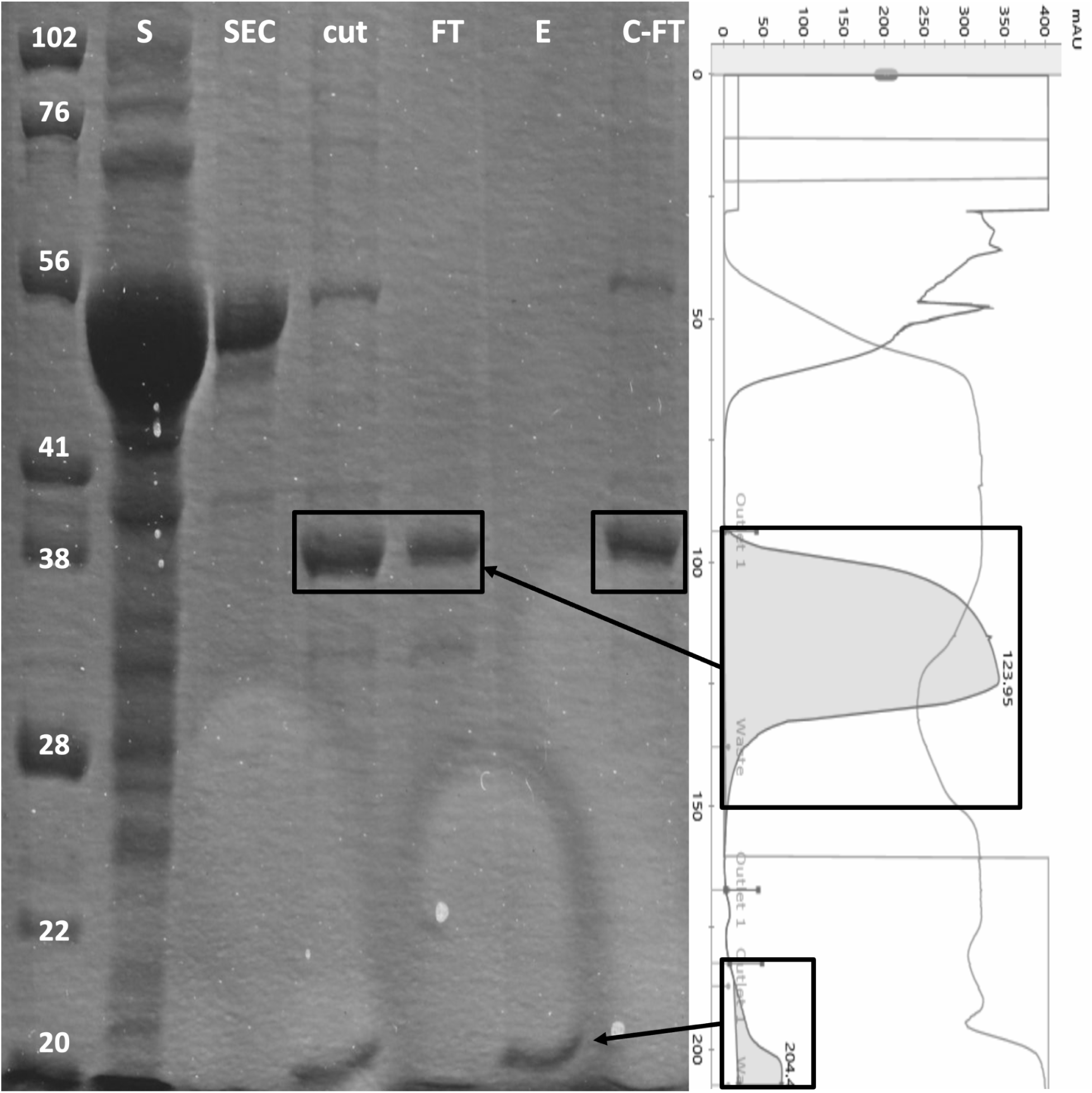
**Purification of HER2-TKD via reverse Ni-NTA affinity after His**₆**-SUMO tag cleavage.** SDS-PAGE and chromatographic analysis of HER2 reverse protein after His₆-SUMO tag cleavage and reverse Ni-NTA affinity purification. A 12% SDS-PAGE gel was used to assess the purification of HER2-TKD (48 kDa) following cleavage of the His₆-SUMO tag (35 kDa) by ULP protease. In the reverse Ni-NTA affinity setup, untagged HER2-TKD does not bind to the Ni-NTA resin and is found in the flow-through, whereas His₆-tagged components—including uncleaved fusion protein, the cleaved SUMO tag, and the His-tagged protease—bind to the column and are later eluted. The gel includes the following samples: a molecular weight marker (lane 1), HER2-TKD supernatant (S) (lane 2), concentrated sample from size-exclusion chromatography (SEC) (lane 3), post-cleavage sample containing tags (cut) (lane 4), flow-through (FT) fraction containing untagged HER2-TKD (lane 5), elution (E) fraction containing His-tagged components (lane 6), and a concentrated flow-through (C-FT) sample enriched in untagged HER2-TKD (lane 7). The chromatogram shown below the gel image depicts the purification profile, including UV absorbance at 280 nm, conductivity, and pH.. The cleavage and purification were carried out at 4 °C for 40 minutes using a 5:1000 ULP:protein ratio.

### 3.3 Crystal Observation

Crystallization studies were carried out on purified HER2-TKD protein. Over 3300 crystallization conditions were tested in each experiment, with Terasaki plates incubated at +4°C and room temperature. Crystals observed after 2-4 weeks of incubation and were regularly checked using microscopy **(Figure 5)**. Our crystallization optimization studies are ongoing, with future efforts focusing on improving buffer compositions, exploring alternative crystallization strategies, and enhancing protein stability to facilitate crystal formation and enable structural analysis.

**Figure 5.**
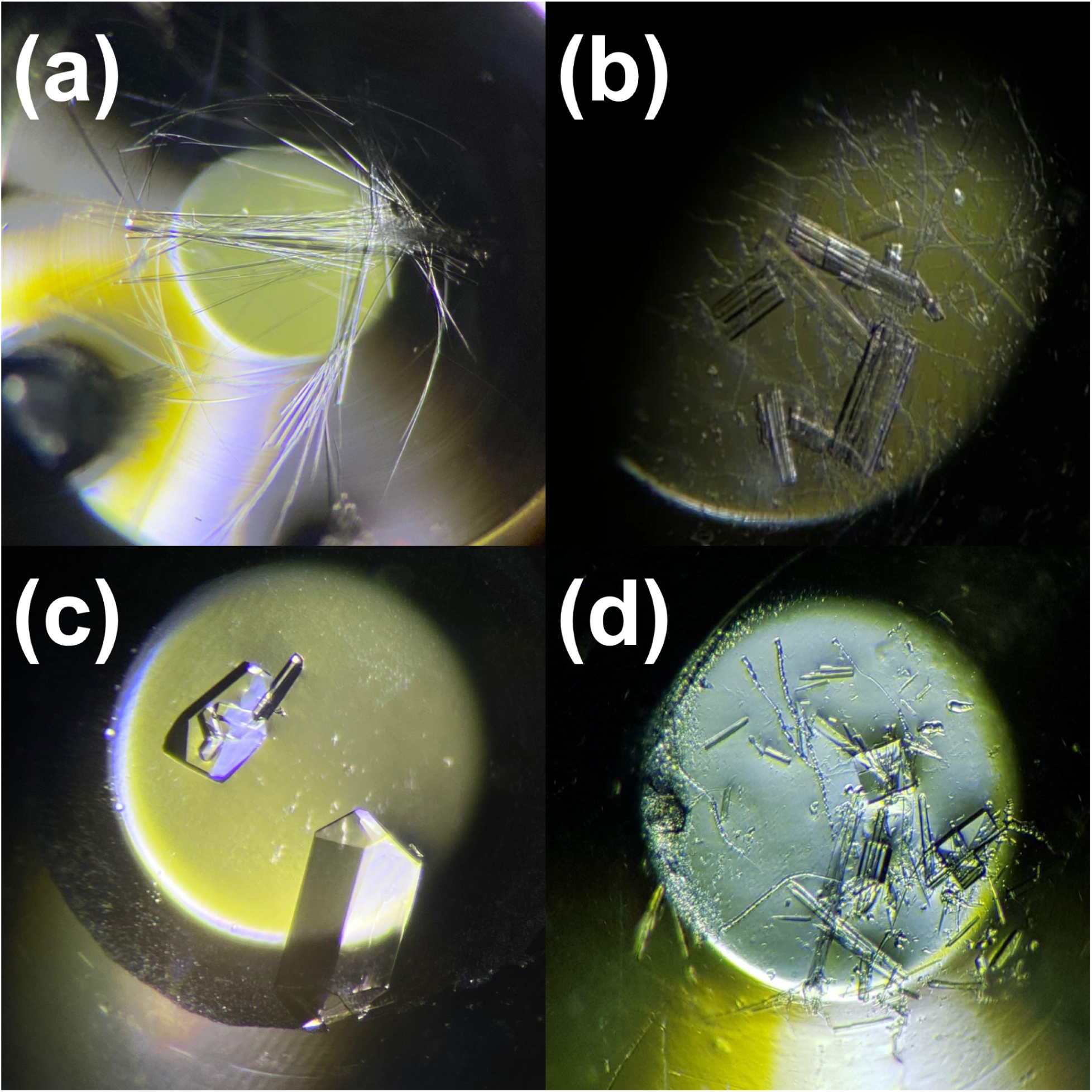
The HER2-TKD crystals obtained through co-crystallization with novel inhibitors. Microscope images of HER2-TKD protein crystals formed after SUMO tag removal and protein concentration. Co-crystallization was carried out in the presence of novel small-molecule inhibitors. **(a)** Needle-like crystals formed in the NRLBD-I #33 condition. **(b)** Crystals obtained in the Pact Premier II #22 condition. **(c)** Block-shaped crystals formed in the Wizard Synergy-II #1 condition. **(d)** Crystals grown in Crystal Screen I #24 condition.

## 4. Discussion

This study represents an important step toward overcoming the challenge of solubilizing HER2-TKD, particularly through the successful use of sarcosyl to eliminate inclusion bodies and recover the protein in a soluble form. While this method significantly improved protein solubility, it also presented a limitation: the inability to perform Ni-NTA affinity purification, most likely due to interference with His-tag binding. Although the specific mechanism remains unclear, this highlights the need for further investigation into the effects of sarcosyl and alternative purification techniques.

Crystallization experiments provided promising results; unfortunately, the majority of the crystals formed were salt crystals rather than protein crystals. Despite this challenge, the findings highlight the inherent complexity of HER2-TKD crystallisation and the difficulties in producing crystals appropriate for high-resolution structural investigation. Thus, further adjustment of crystallization conditions is crucial for enhancing crystal quality and enabling comprehensive structural characterisation.

Overall, our results are important for production and purification strategies for receptor tyrosine kinases such as HER2, particularly by demonstrating an effective method to recover soluble HER2-TKD from inclusion bodies. By expressing HER2-TKD in *E. coli*, we established a cost-effective and efficient production system, making it accessible for structural and functional studies.

To date, numerous small molecules have been investigated as EGFR inhibitors, with several showing promising activity against NSCLC. Our research group has particularly concentrated on developing pyrazoline–thiazole hybrid compounds due to their potential anti-NSCLC properties and EGFR inhibitory effects [11–13]. One such compound, Compound I [11] **(Figure 6)**, demonstrated an IC₅₀ of 10.76 μM for NSCLC inhibition, performing better than erlotinib (IC₅₀ = 22.35 μM). It also showed strong EGFR inhibitory activity with an IC₅₀ of 4.34 μM. Further, Compound II, a naphthyl-substituted pyrazoline-thiazole derivative [12], showed enhanced anti-NSCLC potential with an IC₅₀ of 9.51 μM, outperforming lapatinib (IC₅₀ = 16.44 μM), and inhibited EGFR by 58.32% at 10 μM. In another study by our team [13], the initial scaffold of these hybrids, B-4 **(Figure 6)**, demonstrated more notable anti-NSCLC activity (IC₅₀ = 20.49 ± 2.71 μM) compared to Compound III, the most active molecule in that series. B-4 also achieved 46% EGFR inhibition at 10 μM concentration.

**Figure 6.**
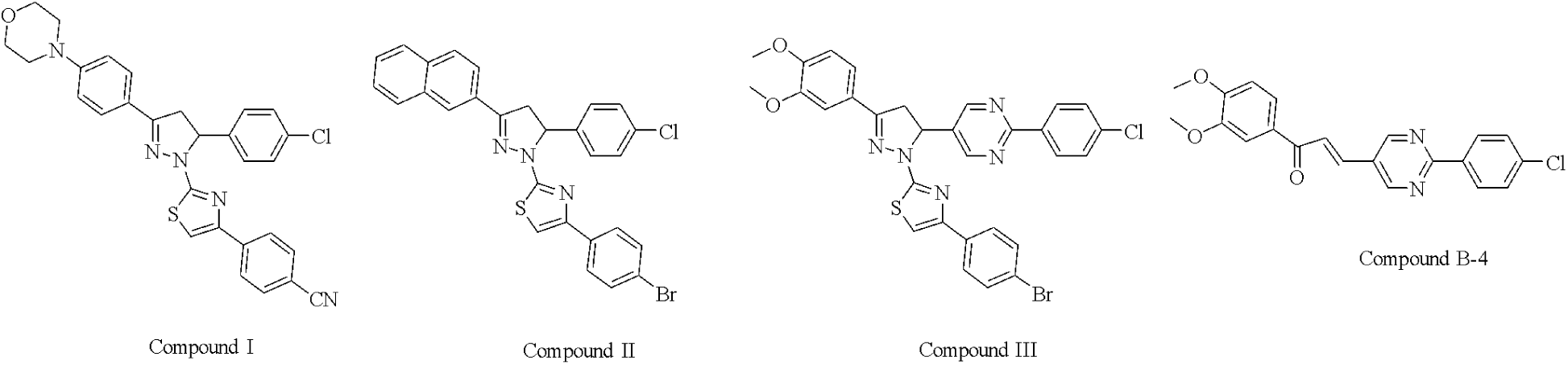
The chemical structures of compounds I, II, III, and B-4

The ongoing development of HER2-specific inhibitors underscores the significance of HER2-TKD as a key target in cancers like breast cancer and NSCLC. In this study, we demonstrated that HER2-TKD can be produced in a soluble form in E. coli in a cost-effective and scalable manner. This achievement not only enables structural studies but also forms the basis of a reliable platform for the screening of potential drug candidates. The successful production, purification, and initial crystallization of HER2-TKD pave the way for obtaining high-resolution structures of HER2-inhibitor complexes, thereby accelerating the discovery of innovative HER2-targeted therapies.

## Acknowledgement

I dedicate this work to my family—especially my mother, Sibel; my father, Hakan; and my dearest husband, Ufuk. I would also like to thank Ahmet Büyükgüngör, Melih Çiğdem, and Sinan Güra for their support, and Belgin Sever for her kindness and generous help.

